# Distributed Clonal Deletion Prevents Autoimmune Disease Progression

**DOI:** 10.64898/2026.05.12.724341

**Authors:** Anna M. Newen, Uzair A. Ansari, Mikala J. Simpson, Fiona Flynn, Sukriti Sharma, Michael Ivanov, Ishaan Antani, Dylan Pfannenstiel, Baktiar Karim, Laura Bassel, Brian Capaldo, Qingrong Chen, Daoud Meerzaman, Indu Raman, Chengsong Zhu, Cornelius Y. Taabazuing, Hamid Kashkar, Christian T. Mayer

## Abstract

Self-reactive B cells are generated during normal development and can acquire increased pathogenicity through activation-induced cytidine deaminase (AID)-mediated diversification following activation. Clonal deletion is thought to eliminate these cells, yet how deletion is distributed across developmental and activation stages to prevent autoimmune disease remains unclear. Here, we show that clonal deletion is enforced through temporally distinct mitochondrial apoptosis (MOMP) checkpoints that differentially regulate autoreactive B cell fate and disease progression. Using conditional Bcl-2 expression to inhibit MOMP either before or after B cell activation, we find that early inhibition permits the survival and maturation of autoreactive B cells after peripheral egress, expanding the pool of cells available for activation. These cells subsequently undergo AID-dependent diversification, producing class-switched IgG autoantibodies with expanded antigen breadth that target a wider range of self-antigens and drive lethal, female-biased autoimmune disease characterized by complement activation and kidney pathology. In contrast, inhibition of MOMP only after activation allows the accumulation of germinal center, switched memory, and plasma cells and promotes autoantibody production, but results in more restricted IgG autoreactivity, limited complement activation and limited tissue damage, and normal survival. Notably, early MOMP inhibition does not expand immature bone marrow B cells, indicating that a major clonal deletion checkpoint operates in the periphery rather than during initial B cell generation. Together, these findings support a Distributed Clonal Deletion Model in which early checkpoints restrict the entry of autoreactive B cells into diversification pathways, while later checkpoints limit the persistence of diversified autoreactive clones, thereby constraining autoimmune disease progression.

**One Sentence Summary:** Distributed clonal deletion prevents autoimmune disease progression by restricting the breadth of autoreactive clones entering immune responses, with early MOMP checkpoints limiting diversification and later checkpoints constraining persistence.

## INTRODUCTION

Systemic autoimmune diseases such as systemic lupus erythematosus (SLE) arise when immune tolerance fails and B cells generate antibodies that target self-tissues. These conditions disproportionately affect women and can lead to chronic inflammation, renal failure, and organ damage. Autoreactive B cells are abundant in healthy individuals (*1, 2*), yet most never produce autoreactive antibodies. Additionally, some individuals produce autoantibodies without developing disease (*3-5*). These three levels of tolerance - autoreactive B cells, autoantibodies, and autoimmune disease - remain incompletely understood. Defining how tolerance mechanisms operate at each level and cooperate to prevent disease progression is essential for understanding why tolerance breaks down in autoimmunity.

B cell development generates a diverse preimmune repertoire through recombination-activating gene (RAG)–mediated V(D)J recombination in the bone marrow. Pro-B cells (Fraction B/C) rearrange immunoglobulin heavy chains and proliferate as large pre-B cells (Fraction C′), followed by light-chain rearrangement in small pre-B cells (Fraction D), and progression to immature B cells (Fraction E) (*6-8*). After leaving the bone marrow, immature B cells mature through transitional stages and enter the follicular (FO) or marginal zone compartments in the spleen (*8-10*). A longstanding but unresolved question is whether immature B cells destined for deletion die within the bone marrow or after peripheral egress (*11-13*), a distinction that has been difficult to resolve because dying cells are rapidly cleared in vivo.

Upon antigen encounter in T cell–dependent immune responses, FO B cells express activation-induced cytidine deaminase (AID). AID mediates somatic hypermutation (SHM), primarily within germinal centers (GCs), which can generate or enhance autoreactivity, and it mediates class-switch recombination (CSR), which frequently occurs early during immune responses and can take place outside GCs (*14-17*). SHM alters antigen specificity, whereas CSR changes antibody isotype and thereby modifies effector properties. Because both RAG and AID can generate or modify autoreactivity, multiple apoptosis checkpoints censor autoreactive B cells (*12, 13, 18-25*), and genetic models that attenuate apoptosis are associated with autoimmune disease (*26-32*). Yet, autoreactivity and autoantibody production do not always progress to disease (*1-5, 33*). Whether checkpoints before and after activation operate independently or cooperatively remains unclear, in part because prior studies of individual tolerance checkpoints have not directly examined how the timing of checkpoint failure across B cell compartments influences progression from autoreactivity to autoimmune disease.

We hypothesized that clonal deletion is distributed across developmental and activation stages and that checkpoint timing determines disease outcomes by controlling which autoreactive B cells undergo AID-dependent diversification. Early checkpoints acting before B cell activation would restrict low-avidity autoreactive clones from undergoing CSR and SHM, whereas late checkpoints acting after activation would limit the persistence of already-diversified clones but act on an already-restricted substrate. Under this model, failure of early checkpoints would broaden the class-switched autoreactive repertoire and drive autoimmune disease progression, whereas failure of late checkpoints would permit autoreactive cell accumulation and autoantibody production without causing organ damage.

To test this hypothesis, we used identical Rosa26^LSL-Bcl2^ alleles driven by distinct Cre recombinases to express Bcl-2 either beginning early in B cell development (Bcl2^Early^) or only after activation (Bcl2^Late^). Bcl-2 inhibits mitochondrial outer membrane permeabilization (MOMP), enabling direct comparison of the consequences of disrupting early versus late MOMP-sensitive tolerance checkpoints. We combined these models with in vivo EdU pulse–chase labeling, comprehensive disease assessment, and autoantibody profiling to define how checkpoint timing regulates B cell progression, AID-dependent diversification, and autoimmune disease outcomes.

## RESULTS

### MOMP inhibition before activation, but not after activation, causes autoimmune disease

We generated mice expressing human Bcl-2 (hBcl-2) and GFP from the Rosa26^LSL-Bcl2^ allele using Mb1^Cre^ or Aicda^Cre^ to inhibit mitochondrial outer membrane permeabilization (MOMP) either from early B cell development (Bcl2^Early^) or only after activation (Bcl2^Late^). Flow cytometry confirmed stage-specific Cre recombination (fig. S1, S2A), and germinal center (GC), switched memory B (swMem), and plasma cells (PC) expressed comparable hBcl-2 levels (fig. S2B, C), enabling direct comparison.

To determine how the timing of MOMP inhibition influences autoimmune disease progression, we monitored disease development in Bcl2^Early^, Bcl2^Late^, and control cohorts. Only Bcl2^Early^ mice developed clinical illness, with reduced survival and a higher incidence of disease in females (Fig. 1A, B). Renal histopathology revealed widespread glomerulonephritis and tubular injury in Bcl2^Early^ mice, characterized by glomerular hypercellularity, capillary wall thickening, and crescent formation, together with inflammatory infiltrates (Fig. 1C). Quantitative scoring confirmed significantly greater pathology in this group than in Bcl2^Late^ and control mice, with more severe changes in females (Fig. 1D–F). Immunoglobulin (Ig) deposition within glomeruli was comparable between Bcl2^Early^ and Bcl2^Late^ mice, whereas cortical Ig deposition was increased in Bcl2^Early^ kidneys (Fig. 1G), indicating that differences in disease severity are not explained by glomerular Ig deposition alone. In contrast, complement C3d deposition was increased only in Bcl2^Early^ mice (Fig. 1H). Serum urea measurements indicated impaired renal function in a subset of Bcl2^Early^ mice (25% of females; 0% of males) (Fig. 1I, J), whereas Bcl2^Late^ mice maintained normal urea levels despite the presence of autoreactive antibodies (Fig. 1I). Together, these findings demonstrate that disruption of early, but not post-activation, MOMP checkpoints drives autoimmune disease progression, whereas post-activation checkpoint failure permits autoreactivity and autoantibody production without progression to organ damage.

**Fig. 1.**
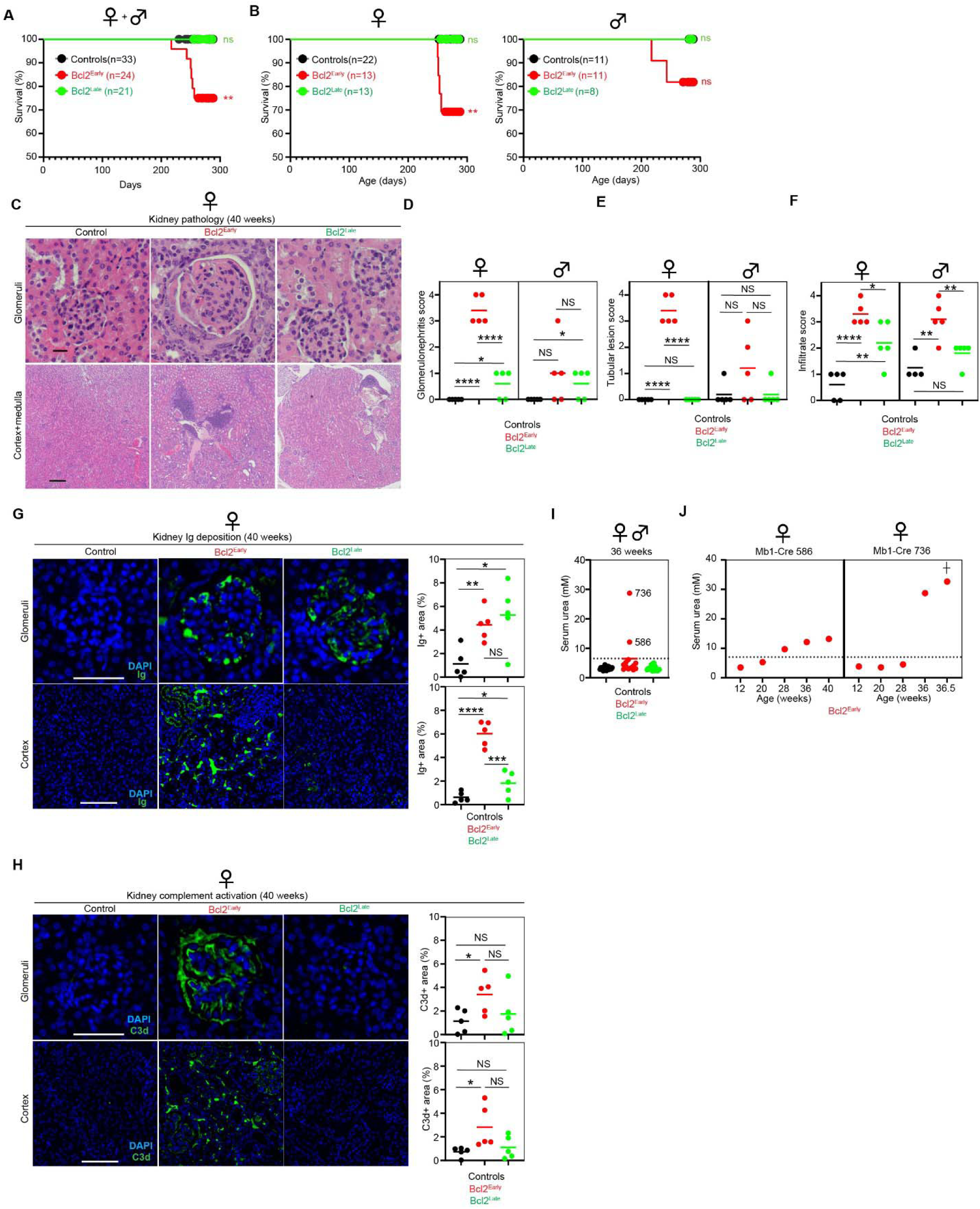
Early, but not post-activation, MOMP inhibition drives autoimmune disease. Cohorts of Bcl2^Early^ (n=24), Bcl2^Late^ (n=21), and Control mice (n=33) were monitored. (**A, B**) Survival analysis of (**A**) all mice combined or (**B**) according to sex. ** p=0.0036 (both sexes), ** p=0.0063 (females), NS=not statistically significant (Log-rank (Mantel-Cox) test). (**C-F**) Analysis of kidney pathology at 40 weeks. **(C)** Representative micrographs depict hematoxylin and eosin-stained kidney sections of females of the indicated genotypes. Top, glomeruli (scale bar: 20µm). Bottom, cortex (scale bar: 200µm). (**D-F**) Pathology scores for (**D**) glomerulonephritis or (**E**) tubular lesion or (**F**) lymphoplasmacytic infiltrates according to sex. **(G, H)** Immunofluorescence analysis of kidney sections. Micrographs to the left depict staining (green) for (**G**) Immunoglobulin (Ig) or (**H**) complement C3d in glomeruli (top, scale bar: 50µm) or cortex (bottom, scale bar: 100µm). DAPI staining is shown in blue. Graphs on the right show Ig^+^ and C3d^+^ area quantification. (**I, J**) Serum urea quantification for (**I**) all mice at 36 weeks or (**J**) longitudinally for two Bcl2^Early^ mice showing urea elevation at 36 weeks of age. Cross indicates terminal disease. Dotted lines represent twice the mean serum urea concentration of control mice, and measurements above this line are considered abnormal. (**D-H**) **** p<0.0001, *** p<0.001, ** p<0.01, * p<0.05, NS=not statistically significant (unpaired two-tailed Student’s t-test). Comparisons relative to controls and between Bcl2^Early^ and Bcl2^Late^ mice are shown. Horizontal bars represent mean values. (**A-J)** Results are from one cohort sequentially recruited from independent litters.

### Bcl-2 expression during development restricts IgG autoantibody breadth

To determine whether checkpoint timing affects autoreactive antibody specificity, we performed autoantigen array analysis (Fig. 2A-C). Bcl2^Early^ mice exhibited a marked expansion of IgG autoreactivity, with 23 significantly increased autoantibodies compared to 5 in Bcl2^Late^ mice relative to controls (Fig. 2A, C and Supplementary Data 1). In contrast, IgM autoreactivity was increased but more comparable between Bcl2^Early^ and Bcl2^Late^ groups (Fig. 2B, C and Supplementary Data 2). Consistent with this, 13 IgG autoantibodies were significantly increased in Bcl2^Early^ compared with Bcl2^Late^ mice, whereas only a single IgM reactivity differed (Fig. 2C), indicating that early checkpoint failure selectively expands class-switched autoreactive breadth rather than overall autoreactivity. Notably, among the most expanded IgG reactivities in Bcl2^Early^ mice were antibodies targeting complement C3 and PCNA, which have been associated with severe nephritis in subsets of patients with systemic lupus erythematosus (*34, 35*). Total autoreactive IgG titers and dsDNA/ssDNA ELISAs were similar between Bcl-2 models (fig. S3A–E), with ELISA confirmation of anti-C3 and anti-PCNA IgG autoantibody differences (fig. S3F, G).

**Fig. 2.**
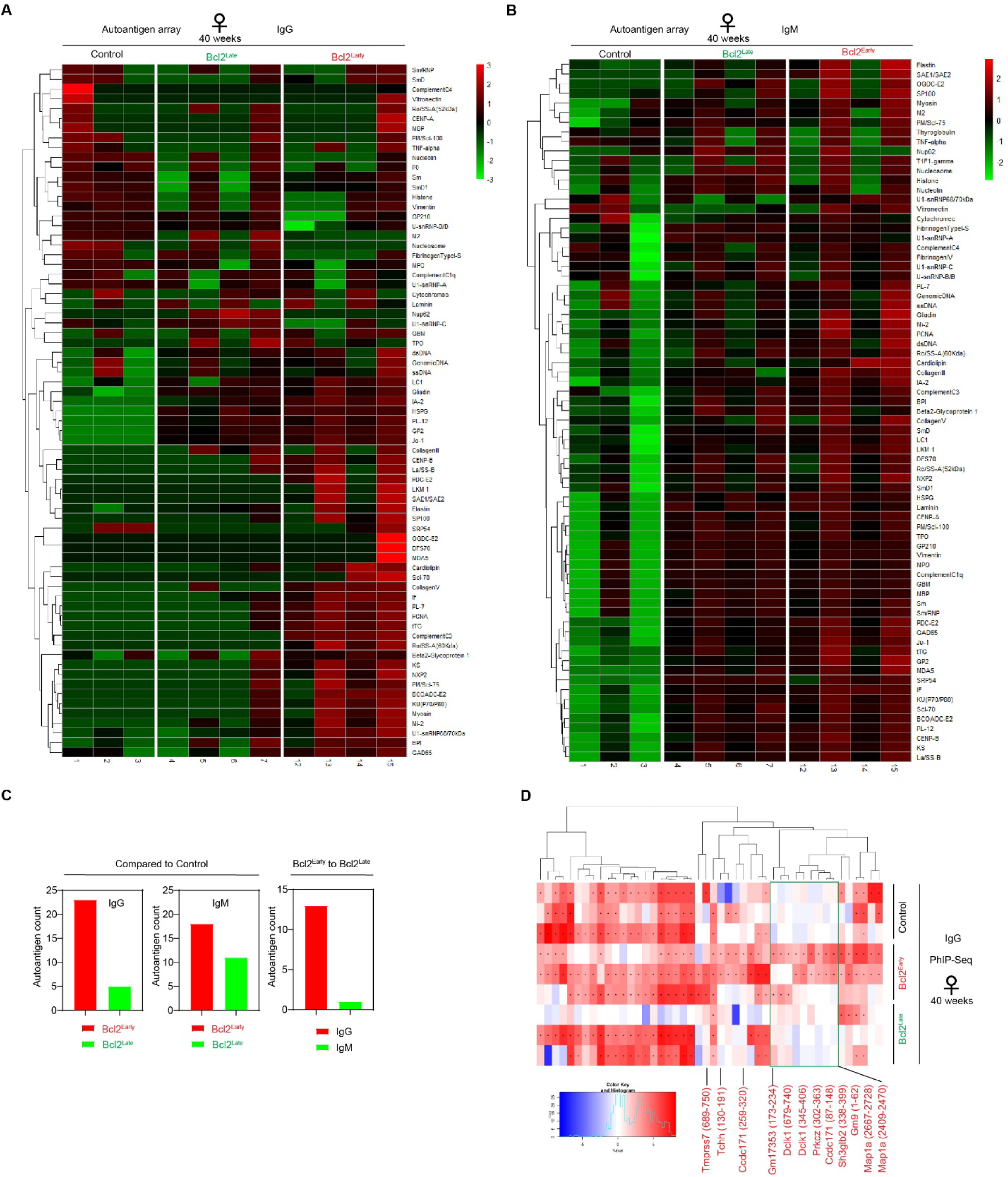
Early MOMP inhibition broadens class-switched IgG autoreactivity. **(A, B)** Autoantigen array heatmaps depict normalized antibody scores (red, high reactivity; green, low reactivity) for (**A**) IgG or (**B**) IgM serum antibody binding to indicated self-antigens. Each column represents one 40-week-old mouse of the indicated genotypes. (**C**) Quantification of autoantibody breadth from autoantigen array data (Supplementary Data 1-2). Left: Number of statistically significant IgG-reactive autoantigens per genotype. Bcl2^Early^ mice show 23 significant IgG autoantigens compared to 5 in Bcl2^Late^ mice relative to controls. Middle: Number of statistically significant IgM-reactive autoantigens per genotype. Bcl2^Early^ mice show 18 significant IgM autoantigens compared to 11 in Bcl2^Late^ mice relative to controls. Right: Number of statistically significant autoantigens in Bcl2^Early^ mice relative to Bcl2^Late^ mice according to isotype. Bcl2^Early^ mice show 13 statistically significant IgG-reactive autoantigens, but only 1 IgM-reactive autoantigen. Early checkpoint failure specifically expands IgG breadth while IgM remains comparable. (**D**) Phage immunoprecipitation sequencing (PhIP-seq) analysis of IgG autoantibodies. Heatmaps display normalized signal intensity (red, high; blue, low). Each row represents one 40-week-old mouse of the indicated genotypes. Dots in the center of a cell depict p<0.05. Only phages yielding significant hits in at least two Bcl2^Early^ mice were included in the heatmap. Antigens encoded by phages yielding no significant hits in control and Bcl2^Late^ mice are labeled in red.

To independently assess autoreactive repertoires, we performed phage immunoprecipitation sequencing (PhIP-seq). Although numerous peptide reactivities were detected across all groups, including controls (fig. S3H), gene set enrichment analysis (GSEA) of peptide-associated genes revealed qualitative differences between genotypes. Bcl2^Early^ mice showed selective enrichment for gene sets related to lipid translocation, intracellular ligand-gated ion channel activity, transcriptional coactivator function, and immune response–associated transcriptional programs (Supplementary Data 3), whereas Bcl2^Late^ mice did not show enrichment of these categories. These findings indicate distinct functional profiles of autoreactive IgG repertoires despite broadly comparable numbers of individual peptide reactivities across groups.

Focusing on peptides enriched in at least two of three Bcl2^Early^ mice but absent from Bcl2^Late^ and control mice identified 12 Bcl2^Early^-specific reactivities, including *Prkcz*, which is associated with autoimmune myopathies in humans (*36*)(Fig. 2D).

Together, these data demonstrate that early checkpoint failure selectively widens class-switched autoreactive IgG breadth, whereas late checkpoint failure permits autoreactivity without broad class-switched diversification.

### AID is required for autoimmune pathology when MOMP is inhibited before activation

Given the increased breadth of class-switched autoantibodies in Bcl2^Early^ mice, we next tested whether AID-dependent processes are required for autoimmune disease in this setting. To do so, we analyzed Bcl2^Early^ mice lacking AID (encoded by Aicda), thereby preventing SHM and CSR. Strikingly, Bcl2^Early^Aicda^-/-^ mice remained free of autoimmune disease and showed no renal injury (Fig. 3A–G), despite exhibiting the same early checkpoint disruption as Bcl2^Early^ mice. Histopathologic evaluation revealed infection-associated mammary gland abscesses (Fig. 3B, C), but no autoimmune pathology (Fig. 3D-G). One Bcl2^Early^ mouse developed B cell lymphoma (Fig. 3B, C). These findings demonstrate that AID-dependent diversification is required for autoimmune disease in the setting of early MOMP-regulated checkpoint failure, and that autoreactive B cell survival without CSR/SHM is insufficient to cause autoimmune organ damage.

**Fig. 3.**
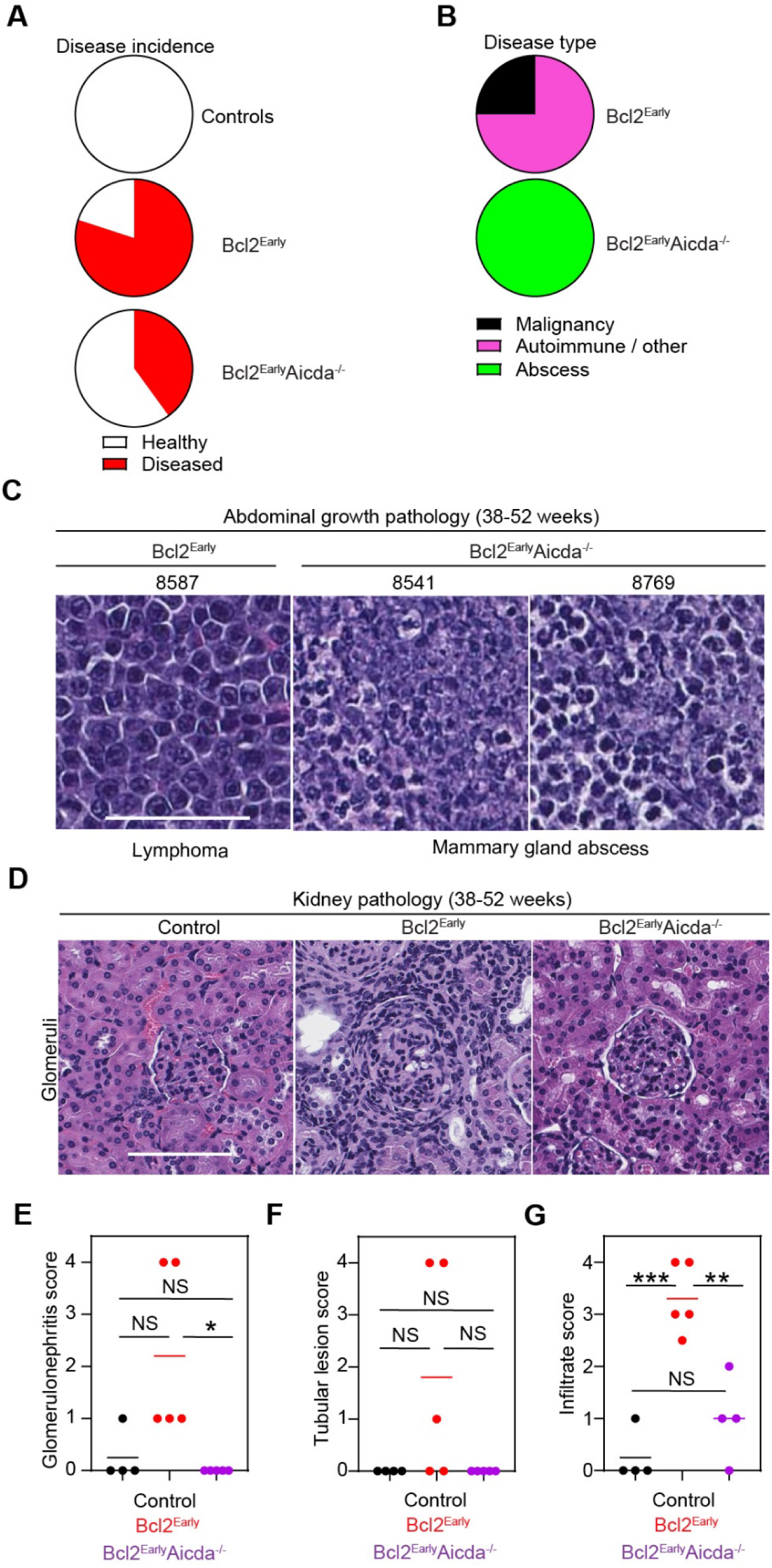
AID is required for autoimmune disease following early MOMP inhibition. Cohorts of female Bcl2^Early^ (n=5), Bcl2^Early^Aicda^-/-^ (n=5), and Control mice (n=4) were monitored. (**A**) Fraction of diseased mice at 52 weeks (white: healthy; red: diseased). (**B**) Type of disease (Autoimmune/other: magenta; B cell malignancy: black; abscess: green). (**C**) Micrographs depict hematoxylin and eosin-stained abdominal mass sections of indicated mice (scale bar: 50µm). Diagnosis is shown below micrographs. **(D)** Representative micrographs depict hematoxylin and eosin-stained kidney sections of 38-52 weeks old females of the indicated genotypes. Glomeruli are shown in the center (scale bar: 100µm). (**E-G**) Pathology scores for (**E**) glomerulonephritis or (**F**) tubular lesion or (**G**) lymphoplasmacytic infiltrates. *** p<0.001, ** p<0.01, * p<0.05, NS=not statistically significant (unpaired two-tailed Student’s t-test). Comparisons relative to controls and between Bcl2^Early^ and Bcl2^Early^Aicda^-/-^ mice are shown. Horizontal bars represent mean values.

### MOMP inhibition before activation expands peripheral B cell populations

To determine how early MOMP checkpoint disruption alters the B cell pool available for activation, we analyzed B cell development in Bcl2^Early^ and Bcl2^Late^ mice (Fig. 4A). Despite Bcl-2 expression beginning at the pro-B stage, bone marrow pro-B, pre-B, and immature B cells were not expanded in Bcl2^Early^ mice, whereas a reduction in large pre-B cells (Fraction C′) was noted, consistent with decreased proliferation (Fig. 4B).

**Fig. 4.**
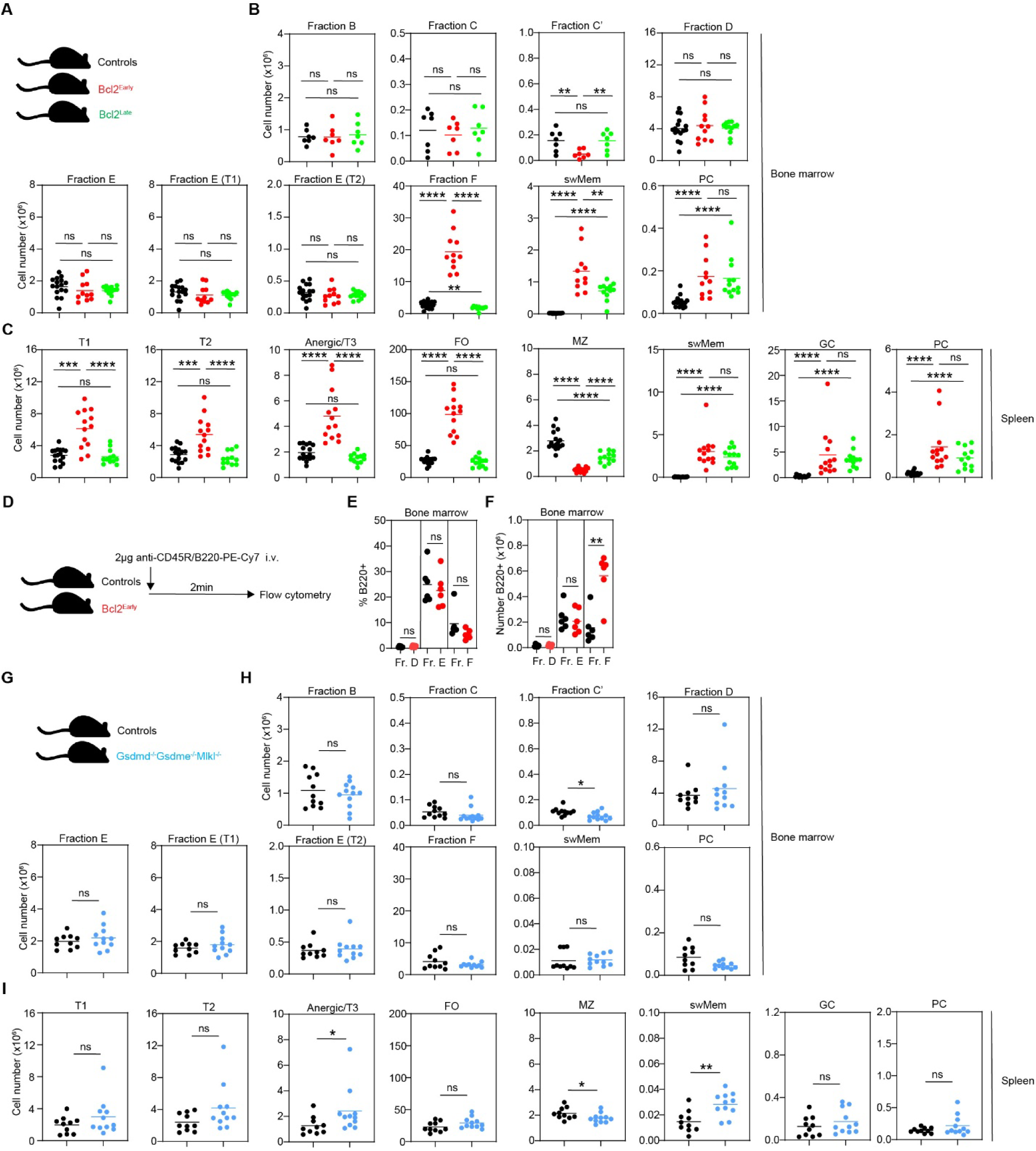
Early MOMP inhibition expands peripheral B cell populations without increasing bone marrow immature B cells. Mice aged 9-13 weeks were analyzed by flow cytometry. (**A**) Genotypes and color coding. (**B, C**) Total numbers of the indicated B cell subsets in (**B**) bone marrow and (**C**) spleen. Data are combined from six independent experiments (Controls, n=16; Bcl2^Early^, n=11 for bone marrow and n=13 for spleen; Bcl2^Late^, n=12). (**D-F**) To assess bone marrow egress, mice aged 14-20 weeks were injected intravenously with anti-CD45R/B220-PE-Cy7 prior to analysis by flow cytometry. (**D**) Experimental scheme. (**E, F**) Quantification of (**E**) the percentage of B220-labeled cells within the indicated B cell subsets and **(F)** the total number of B220-labeled B cell subsets, indicating cells exposed to the bone marrow sinusoids. Data are combined from three independent experiments (Controls, n=6; Bcl2^Early^, n=6). (**G-I**) To test the contribution of non-apoptotic programmed cell death, mice aged 7-17 weeks were analyzed by flow cytometry. (**G**) Genotypes and color coding. (**H, I**) Total numbers of the indicated B cell subsets in (**H**) bone marrow and (**I**) spleen of Gsdmd^-/-^Gsdme^-/-^Mlkl^-/-^ mice and controls. Data are combined from five independent experiments (Controls, n=10; Gsdmd^-/-^Gsdme^-/-^Mlkl^-/-^, n=11). **** p<0.0001, *** p<0.001, ** p<0.01, * p <0.05, NS=not statistically significant (two-tailed Mann-Whitney test for panels B, C, H and I; two-tailed unpaired Student’s t-test for panels E and F). Horizontal bars represent mean values. Abbreviations: swMem, class-switched memory B cells; PC, plasma cells; T1, transitional 1 B cells; T2, transitional 2 B cells; Anergic/T3, anergic B cells; FO, mature follicular B cells; MZ, marginal zone B cells; GC, germinal center B cells.

In contrast, Bcl2^Early^ mice displayed significant expansion of splenic transitional (T1, T2), anergic (T3), and follicular (FO) B cells, as well as increased recirculating mature B cells in the bone marrow (Fig. 4B, C). Bcl2^Late^ mice resembled controls across these compartments (Fig. 4B, C). Both Bcl2^Early^ and Bcl2^Late^ mice accumulated post-activation germinal center (GC) B cells, switched memory B cells (swMem), and plasma cells (PC). These findings indicate that early, but not post-activation, MOMP inhibition selectively expands peripheral immature and mature B cell populations.

### Immature B cell deletion occurs predominantly after bone marrow egress

The absence of immature B cell expansion in Bcl2^Early^ mice suggested either limited apoptosis at this stage or accelerated marrow egress. To test the latter, we performed vascular labeling of bone marrow sinusoids (*11*)(Fig. 4D). Immature B cells labeled by intravenously injected anti-CD45R/B220-PE-Cy7 did not increase in proportion or number in Bcl2^Early^ mice relative to controls (Fig. 4E, F), excluding enhanced marrow egress as a cause of the normal immature pool. Increased numbers of labeled mature B cells (Fraction F) were detected, reflecting expansion and increased bone marrow egress of this compartment (Fig. 4B, E, F).

To determine whether non-apoptotic programmed cell death contributes to immature B cell loss, we analyzed Gsdmd^-/-^Gsdme^-/-^Mlkl^-/-^ triple-deficient mice, which are unable to undergo pyroptosis and necroptosis (Fig. 4G). Bone marrow and splenic B cell subsets were comparable to controls with only modest changes in Fraction C′, T3, and swMem cells (Fig. 4H, I), while macrophages resisted pyroptotic and necroptotic stimuli (fig. S4A). These findings rule out pyroptosis and necroptosis, supporting a MOMP-sensitive peripheral deletion mechanism.

### Transitional B cells are the first peripheral stage regulated by MOMP

To define how MOMP inhibition influences developmental progression, we performed EdU pulse–chase labeling (Fig. 5A). In control mice, EdU labeled Fraction C′ within two hours and appeared in Fraction E, T1, T2, and T3 by day 3, with labeled transitional and anergic populations contracting by day 7 and labeled FO and Fraction F cells persisting (Fig. 5B, C).

**Fig. 5.**
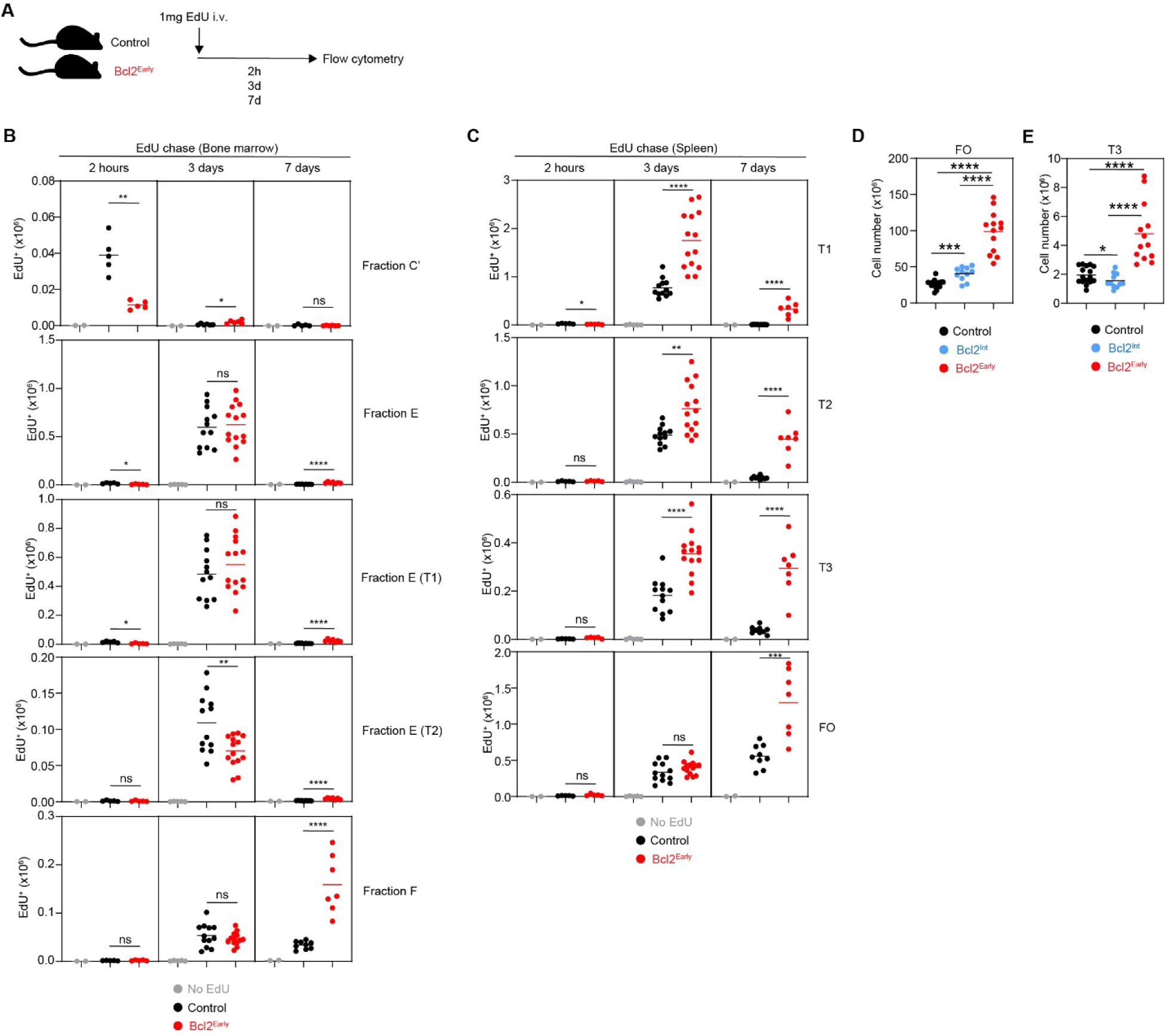
Transitional B cell selection limits follicular B cell expansion. Mice aged 8-18 weeks were injected intravenously with 1mg EdU and analyzed by flow cytometry at the indicated chase times. (**A**) Experimental schematic. (**B, C**) Total numbers of the indicated EdU^+^ B cell subsets in (**B**) bone marrow and (**C**) spleen. Data are combined from two (2h, 7d) or five (3d) independent experiments (2h: Controls, n=5; Bcl2^Early^, n=5; 3d: Controls, n=12; Bcl2^Early^, n=14; 7d: Controls, n=9; Bcl2^Early^, n=7; Fr. C’, n=5 per genotype for all time points). (**D, E**) Steady-state total numbers of (**D**) follicular (FO) and (**E**) anergic T3 B cells in the spleen, analyzed by flow cytometry in mice aged 9–13 weeks. Bcl2^Int^ mice are combined from four independent experiments (n = 11) run in parallel with the same Control and Bcl2^Early^ cohorts shown in Fig. 4. **** p<0.0001, *** p<0.001, ** p<0.01, * p <0.05, NS=not statistically significant (two-tailed unpaired Student’s t-test). Horizontal bars represent mean values. Abbreviations: T1, transitional 1 B cells; T2, transitional 2 B cells; T3, anergic B cells; FO, mature follicular B cells.

In Bcl2^Early^ mice, EdU labeling of Fraction C′ cells was reduced at two hours (Fig. 5B), consistent with decreased proliferation. Fraction E labeling on day 3 was unchanged, whereas splenic EdU T1, T2, and T3 cells were significantly increased at day 3 and remained elevated at day 7 (Fig. 5C). Early FO cells appeared similarly at day 3; however, by day 7, FO and Fraction F B cells were significantly expanded in Bcl2^Early^ mice (Fig. 5C). These findings identify transitional and anergic B cells as the first peripheral populations whose survival is limited by MOMP.

### FO-intrinsic MOMP inhibition is insufficient to account for peripheral expansion

To assess the contribution of FO-intrinsic MOMP inhibition, we examined CD21^Cre^Rosa26^LSL-Bcl2^ (Bcl2^Int^) mice, in which recombination is efficient in FO B cells and partial in T2 and anergic T3 cells (*37*)(fig. S4B). FO B cells were increased in Bcl2^Int^ mice relative to controls but remained substantially fewer than in Bcl2^Early^ mice (Fig. 5D). Transitional and anergic T3 B cell numbers in Bcl2^Int^ mice did not differ from controls (Fig. 5E and fig S4C, D). These findings indicate that FO-intrinsic MOMP inhibition contributes to but is insufficient to account for the Bcl2^Early^ phenotype, highlighting the importance of early peripheral checkpoints acting at transitional stages.

Together, these findings suggest that early MOMP checkpoint disruption permits the survival of autoreactive B cells after peripheral egress, leading to expansion of transitional and follicular B cell populations, increased class-switched IgG breadth, and AID-dependent autoimmune disease, whereas post-activation checkpoint failure permits autoreactivity without progression to pathology.

## DISCUSSION

This study demonstrates that the timing of MOMP-sensitive checkpoints determines whether autoreactive B cells progress to autoimmune disease. Inhibition of MOMP before B cell activation permits the survival and expansion of peripheral B cell populations, leading to increased class-switched autoreactive IgG breadth and severe, AID-dependent autoimmune pathology. In contrast, MOMP inhibition restricted to activated and post-germinal center stages allows autoreactive B cell accumulation and autoantibody production but does not result in organ damage. These findings provide a mechanistic explanation for how autoreactivity and autoantibodies can be present in the absence of clinical disease (*1, 3, 4*).

Our results extend prior work on apoptosis checkpoints in B cells by directly comparing the consequences of disrupting early versus late MOMP-sensitive stages. Previous studies demonstrated apoptosis in developing and germinal center B cells (*11, 38-40*) and identified a post-germinal center checkpoint that limits autoreactive plasma cell and memory B cell survival (*33, 41*). However, how checkpoint timing influences progression from autoreactivity to autoimmune disease remained unresolved. Here, we show that early checkpoint disruption uniquely permits disease progression, whereas late checkpoint disruption alone is insufficient in our model, establishing that these checkpoints make non-equivalent contributions to immune tolerance.

Mechanistically, early MOMP inhibition expands transitional and follicular B cell populations without altering immature bone marrow subsets, indicating that a major clonal deletion checkpoint operates after peripheral egress. This extends prior apoptotic cell quantification (*11*) by defining functional consequences through genetic loss-of-function approaches. EdU tracing further identifies transitional B cells as the first stage at which survival is MOMP-limited, with downstream expansion of follicular populations. These findings suggest that early peripheral checkpoints constrain the size and composition of the B cell pool entering immune responses, whereas later checkpoints primarily regulate the persistence of activated clones.

The selective expansion of class-switched autoreactive IgG breadth in Bcl2^Early^ mice provides a functional link between checkpoint timing and disease outcome. Despite comparable IgM autoreactivity and similar total autoreactive IgG titers, early checkpoint disruption broadened IgG specificity, including reactivities associated with severe SLE manifestations (*34, 35*). In contrast, Bcl2^Late^ mice exhibited autoreactive antibodies without comparable IgG diversification or pathology. Together with the complete protection observed in AID-deficient Bcl2^Early^ mice, these findings demonstrate that AID-dependent diversification is required for disease and that survival of autoreactive B cells alone is insufficient to drive pathology in this model. AID has also been implicated in B cell tolerance (*42-45*), and AID deficiency has been reported to either attenuate (*46, 47*) or exacerbate (*48, 49*) autoimmune pathology depending on the model. In our system, AID deficiency completely prevented disease despite early checkpoint disruption, indicating that the pathogenic effects of AID-dependent diversification outweigh potential tolerance-promoting functions under these conditions. These findings suggest that the impact of AID on autoimmunity is context-dependent and shaped by the availability of autoreactive precursors.

Notably, immunoglobulin deposition within glomeruli was observed in both Bcl2^Early^ and Bcl2^Late^ mice, yet complement activation and renal pathology occurred only in Bcl2^Early^ animals. This dissociation suggests qualitative differences in the autoreactive antibody response rather than differences in total antibody deposition. One possibility is that the antigenic targets differ, as Bcl2^Early^ mice exhibit broader IgG reactivity, including antibodies such as anti-C3 that have been shown to enhance complement activation by interfering with regulatory mechanisms or stabilizing C3 convertases (*50, 51*). Alternatively, differences in antibody isotype may influence complement fixation and pathogenic potential. These considerations indicate that the composition of the autoreactive repertoire, rather than the presence of immune complexes alone, is a key determinant of downstream tissue injury.

These observations support a model in which clonal deletion is distributed across pre-activation and post-activation stages. Because receptor editing efficiently removes high-avidity autoreactive B cells in the bone marrow with minimal reliance on deletion (*11, 52, 53*), we propose that early peripheral checkpoints limit the entry of low-affinity or low-avidity autoreactive B cells into immune responses, thereby restricting the breadth of class-switched autoreactive repertoires, whereas later checkpoints constrain the persistence and expansion of autoreactive clones once activated (Fig. 6). Failure of early checkpoints expands the pool of cells available for diversification and enables pathogenic autoantibody responses, while failure of late checkpoints permits autoreactivity without disease progression (Fig. 6). This framework, which we term the **Distributed Clonal Deletion Model**, provides a unifying explanation for how substantial autoreactivity can be tolerated in healthy individuals yet lead to disease when early tolerance mechanisms fail.

**Fig. 6.**
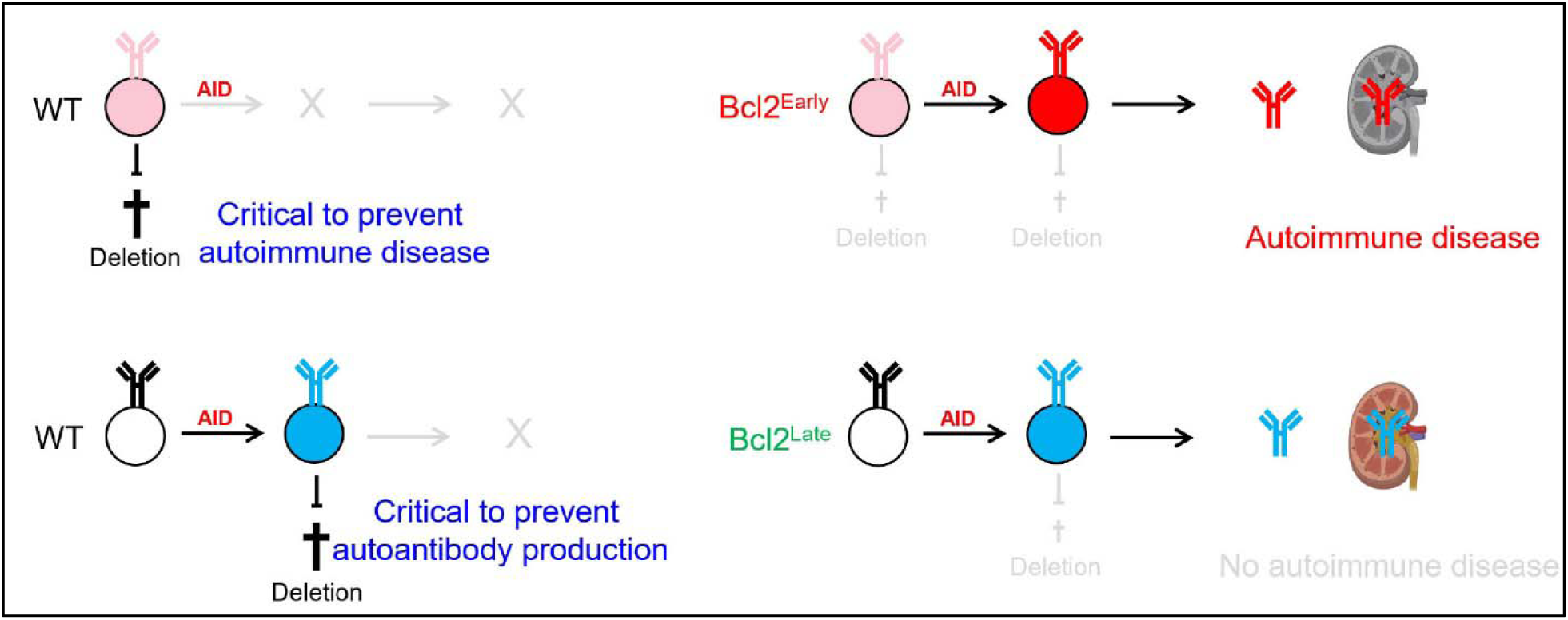
Distributed Clonal Deletion Model of B Cell Tolerance. Schematic illustrating the regulation of B cell tolerance by apoptosis. Receptor editing in the bone marrow efficiently eliminates high-affinity or high-avidity autoreactive immature B cells, independent of genotype, allowing the release of low-affinity or low-avidity autoreactive B cells (light red) and non-autoreactive B cells (white) into the periphery. In wild-type mice, these cells undergo apoptosis prior to activation, representing a key tolerance checkpoint that prevents autoimmunity (top left). In addition, autoreactive B cells generated during immune responses through somatic hypermutation (blue) are deleted after activation, limiting autoantibody production (bottom left). Early Bcl-2 expression disrupts this process by preventing the deletion of low-affinity or low-avidity autoreactive B cells, enabling their survival, entry into the follicular compartment, and participation in immune responses. These cells undergo AID-mediated diversification, generating class-switched, high-affinity autoreactive B cells that produce autoantibodies. The resulting immune complexes deposit in the kidney, activate complement, and drive severe renal pathology and loss of function (top right). In contrast, late Bcl-2 expression does not affect pre-activation tolerance checkpoints but promotes the survival of autoreactive B cells generated by somatic hypermutation. This leads to class-switched autoantibody production and immune complex deposition in the kidney, but with minimal complement activation, minimal pathology, and no significant loss of renal function (bottom right). This model is conceptual and not quantitative.

These findings have implications for human autoimmune disease, where defects in early tolerance checkpoints or altered survival of autoreactive precursors may expand the repertoire available for pathogenic diversification. Targeting pathways that promote autoreactive B cell deletion or limit class-switched autoantibody diversification and complement activation, rather than global B cell depletion, may therefore represent a strategy to reduce autoimmune pathology while preserving immune competence.

## Limitations of the study

This study has several limitations. Bcl-2 inhibits MOMP but may also affect additional cellular processes, and thus our findings define MOMP-sensitive checkpoints rather than apoptosis-specific mechanisms. In addition, although AID dependence is demonstrated, the relative contributions of SHM and CSR were not resolved.

## MATERIALS AND METHODS

### Study Design

This study compared the effect of MOMP inhibition before or after B cell activation on B cell accumulation, autoantibody formation, and autoimmune pathology. Experiments were designed to analyze B cell subset composition, developmental kinetics, serological, and pathological outcomes in Bcl2^Early^, Bcl2^Late^, and control mice. Both male and female mice were analyzed. Sample sizes were based on prior studies demonstrating sufficient statistical power to detect differences in B cell subset frequencies and autoantibody reactivity. No animals or data points were excluded.

### Animals

Aicda^Cre^ (*54*), C57Bl/6J, Gsdmd^-/-^ (*55*), Gsdme^-/-^ (*56*) and Mb1^Cre^ (*57*) mice were from Jackson Laboratories. Aicda^-/-^ (*58*) mice were provided by Dr. Michel Nussenzweig (The Rockefeller University). CD21^Cre^ (*37*) mice by Dr. Jagan Muppidi (NCI). Mlkl^-/-^ (*59*) mice by Dr. James Murphy (WEHI) and Dr. Alan Sher (NIAID), and Rosa26^LSL-Bcl2-IRES-GFP^ mice (*60*) by Dr. Hamid Kashkar (University of Cologne). All mice were on C57Bl/6 background. Some mice were injected intravenously with 1mg 5-ethynyl-2’-deoxyuridine (EdU) (ThermoFisher, A10044) 2h, 3d or 7d prior to analysis, or with 2µg anti-CD45R/B220-PE-Cy7 (ThermoFisher, 25-0452-82) 2min prior to analysis. Mice were maintained under specific pathogen-free conditions at NCI (Frederick and Bethesda). All procedures were approved by the NCI Animal Care and Use Committee (ACUC) and conformed with federal regulatory requirements and standards. The intramural NIH ACU program is accredited by AAALAC International.

### Pathology

Tissues were fixed in buffered 10% formalin (Azer scientific, PFNBF-120) for 48 hours at room temperature and stored in 70% ethanol prior to paraffin embedding. Sections were stained with hematoxylin & eosin and evaluated by a board-certified veterinary pathologist. All slides were evaluated in a blinded fashion using an Olympus BX46 microscope at 200× magnification. Images were captured using a Nikon DS-Ri2 camera. Two pathologists, blinded to genotype, graded glomerulopathy on a scale from 0 to 4 based on the extent and severity of glomerular changes. Grade 0 represented normal glomeruli, without abnormalities. Grade 1 indicated minimal hypercellularity involving less than 25% of glomeruli. Grade 2 was defined by mild hypercellularity affecting 25-50% of the glomerular tufts. Grade 3 denoted moderate hypercellularity in greater than 50 to 75% of glomeruli, with or without crescent formation and thickening of basement membrane. Grade 4 represented marked hypercellularity in greater than 75% of glomeruli, accompanied by crescent formation and thickening of basement membrane. Cortical tubular lesions were graded on a scale from 0 to 4 based on the extent of cortical involvement. Grade 0 (normal) indicated normal tubules without any pathological changes. Grade 1 (minimal) represented tubular lesions, including degeneration and/or regeneration, affecting less than 10% of the cortical area. Grade 2 (mild) involved tubular lesions affecting more than 10% but less than 25% of the cortical tubules. Grade 3 (moderate) indicated lesions involving more than 25% but less than 50% of the cortical tubules. Grade 4 (severe) denoted tubular lesions affecting more than 50% of the cortex. Additionally, lymphoplasmacytic infiltrates in the kidney were semi-quantitatively graded on a scale from 0 to 4, where 0 was normal, 1 was minimal, 2 was mild, 3 was moderate, and 4 was severe.

### Enzyme-linked immunosorbent assay (ELISA)

Serum IgG autoantibodies were measured by ELISA as described (*41*) using peroxidase-conjugated goat anti-mouse IgG Fc (Jackson ImmunoResearch, 115-035-164). Self-antigens included dsDNA (Sigma, D4522), single stranded DNA (ssDNA, prepared from dsDNA), mouse C3 (Complement Technology, M113), and human PCNA (Novus Biologicals, NBC1-18428). Serum was tested at a 1:640 dilution and at three four-fold dilutions (dsDNA, ssDNA), or at a 1:160 dilution (C3, PCNA). Monoclonal control antibodies (mGO53, non-reactive; ED38, highly polyreactive) with mouse IgG1 constant regions were tested at 4µg/ml and at three four-fold dilutions. Absorbance at 405nm was measured with a SpectraMax iD3 Multi-Mode Microplate Reader (Molecular Devices). A_405_ values were corrected for PBS-only signal and area under the curve (AUC) values were calculated using GraphPad Prism.

### Kidney function

Serum urea concentrations were measured with the Urea Nitrogen (BUN) Colorimetric Detection Kit (Thermo Fisher Scientific, EIABUN) according to the manufacturer’s instructions. Measurements were performed with a SpectraMax iD3 Multi-Mode Microplate Reader (Molecular Devices).

### Flow cytometry

Flow cytometry was performed as described (*11*). EdU was detected using the Click-iT™ Plus EdU Pacific Blue™ Flow Cytometry Assay Kit (Cat. C10636, Thermo Fisher). For antibody details see Supplementary Table 1.

### FLOWMIST assay for autoreactivity

The flow cytometry assay was performed as described (*11*). Serum was diluted 1:160 and autoreactive antibodies detected with 1µg/ml AlexaFluor647-conjugated F(ab’)2 goat anti-mouse IgG Fc (Jackson ImmunoResearch, 115-606-071). End titers were the last dilution producing a mean fluorescence intensity (MFI) ratio >3 over negative control serum.

### Autoantigen array

Autoantigen arrays were performed as described (*33*) with the following differences: Serum was tested at 1:640 dilution in PBS. Autoantibodies were detected with Cy3-conjugated goat anti-mouse IgG(H+L) (Invitrogen, Cat. A10521) and with AlexaFluor633-conjugated goat anti-mouse IgM (Invitrogen, Cat. No. A21046) secondary antibodies.

### Phage Immunoprecipitation Sequencing (PhIP-Seq)

PhIP-Seq was performed by CDI Labs as described (*61, 62*). Counts provided by the vendor were analyzed using edgeR (*63*). Modifications to the edgeR workflow were followed as described (*64*). Gene set enrichment analyses and over representation analyses were performed using the clusterProfiler package with the msigdbr package (https://CRAN.R-project.org/package=msigdbr) used for the gene set database (*65*).

### Complement staining

Staining was performed on 5μm FFPE sections using a manual benchtop method. Antigen retrieval was performed with EDTA buffer for 10mins at 100°C. Nonspecific binding was blocked with an incubation of 2% normal donkey serum for 20 mins. This was followed by an overnight incubation at 4°C with the C3d primary antibody (R&D Systems, AF2655) at a 1:250 dilution. Antibody detection was accomplished by incubation with Donkey anti-Goat AlexaFluor 594 (Invitrogen, A11058), followed by incubation in DAPI. Slides were digitally imaged at 20x using a Leica Aperio FL fluorescence digital scanner.

### Immunoglobulin (Ig) staining

FFPE blocks were sectioned at 5μm in preparation for IgM, IgG, IgA IHC. Staining was performed using a Leica Bond RX autostainer (Leica Biosystems). Antigen retrieval was performed using EDTA for 20 mins at 100°C on the Bond autostainer. Endogenous biotin was blocked using the Avidin-Biotin blocking kit (Vector Labs, SP-2001) per the manufacturer’s instructions. The primary antibody for IgM, IgG, and IgA (Southern Biotechnology, 1012-08) was diluted 1:100, with a 30-minute incubation time. Antibody detection was accomplished using AlexaFluor 660 conjugated Streptavidin (Invitrogen, S21377) followed by DAPI counterstain. Slides were digitally imaged at 20x using a Leica Aperio FL fluorescence digital scanner.

### Image analysis

Image analysis was conducted using the Area Quantification FL v2.3.3 algorithm in HALO (Indica Labs, Albuquerque, NM). Analyses were conducted for immunoglobulin / complement staining within the cortex and glomeruli to quantify the percentage of positive area.

### Bone marrow derived macrophage stimulation

Bone marrow derived macrophages (BMDM) were differentiated in RPMI1640 with 10% heat-inactivated fetal bovine serum, penicillin/streptomycin, and recombinant Fc-tagged human M-CSF (amino acids 33 to 190; produced and purified in-house) for 6 days, then primed with 1μg/ml Pam3CSK4 (InvivoGen, tlrl-pms) for 5h. To induce pyroptosis, BMDM were stimulated with 10μM nigericin (Invivogen, tlrl-nig) for 1h. To induce necroptosis, BMDM were stimulated with 100ng/ml TNFα (Sigma, T7539-10UG), 500nM birinapant (Apexbio, A4219) and 20μM Z-VAD-FMK (Tocris, 2163) for 16h.

### Statistical analysis

Statistical significance was determined using GraphPad Prism software. Data were first evaluated for normal distribution using the Anderson-Darling test, D’Agostino & Pearson test, Shapiro-Wilk test and Kolmogorov-Smirnov test. If any test reported that N is too small for evaluation or if data were normally distributed, a two-tailed unpaired t-test was used. If data did not pass normality tests, Mann-Whitney test was used. Survival curves were evaluated using the Log-rank (Mantel-Cox) test. Test results are indicated in the Figures and Figure legends.

## Supporting information

Newen et al. Supplementary Information

## Supplementary Materials

fig S1. Gating strategy for B cell subset identification

fig S2. Validation of GFP and hBcl-2 expression in Bcl2^Early^ and Bcl2^Late^ mice fig S3. Serum IgG autoantibody reactivity

fig S4. Control experiments and validation of Bcl2^Int^ mice

Supplementary Table 1. Details for monoclonal antibodies used in this study Supplementary Data 1. IgG autoantigen array data

Supplementary Data 2. IgM autoantigen array data

Supplementary Data 3. Gene-set enrichment analysis of IgG autoreactivity assessed by phage immunoprecipitation sequencing

During the preparation of this work the authors used Claude and ChatGPT provided by the U.S. Department of Health and Human Services to optimize language. After using these tools, the authors reviewed and edited the content as needed and take full responsibility for the content of the published article.

## Acknowledgments

We thank all members of the Experimental Immunology Branch, and Drs. Ben Afzali, Avinash Bhandoola, Didier Portilla and Jagan Muppidi for discussions and advice; Jeffrey Chiang and Jie Mu for technical support; Assiatu Crossman, Kheem Bisht, Don Plugge, William Hajjar, and Tengfei Zhang for flow cytometry support; all staff at the NCI Frederick and NCI Bethesda animal facilities for their critical help, particularly Jennifer Wise.

## Disclaimer

This research was supported by the Intramural Research Program of the National Institutes of Health (NIH). The contributions of the NIH author(s) were made as part of their official duties as NIH federal employees, are in compliance with agency policy requirements, and are considered Works of the United States Government. However, the findings and conclusions presented in this paper are those of the author(s) and do not necessarily reflect the views of the NIH or the U.S. Department of Health and Human Services.

## Author contributions

Conceptualization: CTM

Methodology: CTM, MJS, CYT, HK

Investigation: CTM, AMN, UAA, MJS, FF, SS, MI, IA, BK, LB, IR, CZ, BC, QC, DM

Visualization: CTM, AMN, UAA, MJS, FF, DP, BK, LB, BC, QC, DM, IR, CZ

Funding acquisition: CTM

Project administration: CTM

Supervision: CTM

Writing – original draft: CTM

Writing – review & editing: CTM, AMN, UAA, MJS, FF, SS, MI, IA, DP, BK, LB, BC, QC, DM, IR, CZ, CYT, HK

## Data and materials availability

All data are available in the main text or the supplementary materials. Autoantigen array data are available at GEO accession GSE296597. PhIP-seq data are available at NCBI accession PRJNA1271546

