## Supplementary material for "Distributed Clonal Deletion Prevents Autoimmune Disease Progression": Newen et al. Supplementary Information

**Supplementary Tables**

| **Antigen** | **Antibody clone** | **Conjugate** | **Supplier** | **Catalog #** | **Dilution** |
| --- | --- | --- | --- | --- | --- |
| CD4 | RM4-5 | Biotin | BioLegend | 100508 | 1:200 |
| CD5 | 53-7.3 | PE-Cy7 | BioLegend | 100622 | 1:200 |
| CD8α | 53-6.7 | Biotin | BD | 553029 | 1:200 |
| CD16/CD32 | 2.4G2 | Unlabeled | In-house | N/A | 1:10 |
| CD19 | 6D5 | AlexaFluor594 | BioLegend | 115552 | 1:800 |
| CD21/CD35 | 7G6 | BUV805 | BD | 741961 | 1:800 |
| CD23 | B3B4 | BV786 | BD | 563988 | 1:200 |
| CD24 | M1/69 | BUV805 | Thermo Fisher | 368-0242-82 | 1:640 |
| CD43 | S7 | PE-Cy7 | BD | 562866 | 1:160 |
| CD43 | S7 | PerCP-Cy5.5 | BD | 562865 | 1:25 |
| CD43 | S7 | BUV805 | BD | 741931 | 1:200 |
| CD45R/B220 | RA3-6B2 | FITC | BD | 553088 | 1:200 |
| CD45R/B220 | RA-3-6B2 | PE | BD | 553090 | 1:80 |
| CD45R/B220 | RA-3-6B2 | AlexaFluor594 | BioLegend | 103254 | 1:400 |
| CD93/AA4.1 | AA4.1 | APC | Thermo Fisher | 17-5892-83 | 1:100 |
| CD95/Fas | Jo2 | R718 | BD | 752226 | 1:3200 |
| CD138 | 281-2 | BV711 | BioLegend | 142519 | 1:8000 |
| CD267/TACI | ebio8F10-3 | PE | eBioscience | 12-5942-81 | 1:200 |
| Bcl-2 | 100 | BV421 | Biolegend | 658709 | 1:20 |
| F4/80 | BM8 | Biotin | Thermo Fisher | 13-4801-85 | 1:200 |
| GL7 | GL7 | Biotin | BioLegend | 144616 | 1:3200 |
| GL7 | GL7 | FITC | BioLegend | 144603 | 1:400 |
| GL7 | GL7 | Pacific Blue | BioLegend | 144614 | 1:3200 |
| IgD | 11-26c.2a | BUV395 | BD | 564274 | 1:100 |
| IgM | RMM-1 | BV421 | BioLegend | 406517 | 1:20 |
| IgM | II/41 | PE-Cy7 | Thermo Fisher | 25-5790-82 | 1:400 or 1:1200 |
| IgM | II/41 | eFluor450 | Invitrogen | 48-5790-82 | 1:160 |
| Ly-6G | 1A8 | Biotin | BioLegend | 127604 | 1:100 |
| Ly-51/BP-1 | 6C3 | PE | BioLegend | 108308 | 1:40 |
| NK1.1 | PK136 | Biotin | BD | 553163 | 1:200 |
| TER-119 | TER-119 | Biotin | Thermo Fisher | 13-5921-82 | 1:400 |

**Supplementary Table 1.** Details for monoclonal antibodies used in this study.

**Supplementary Figures**

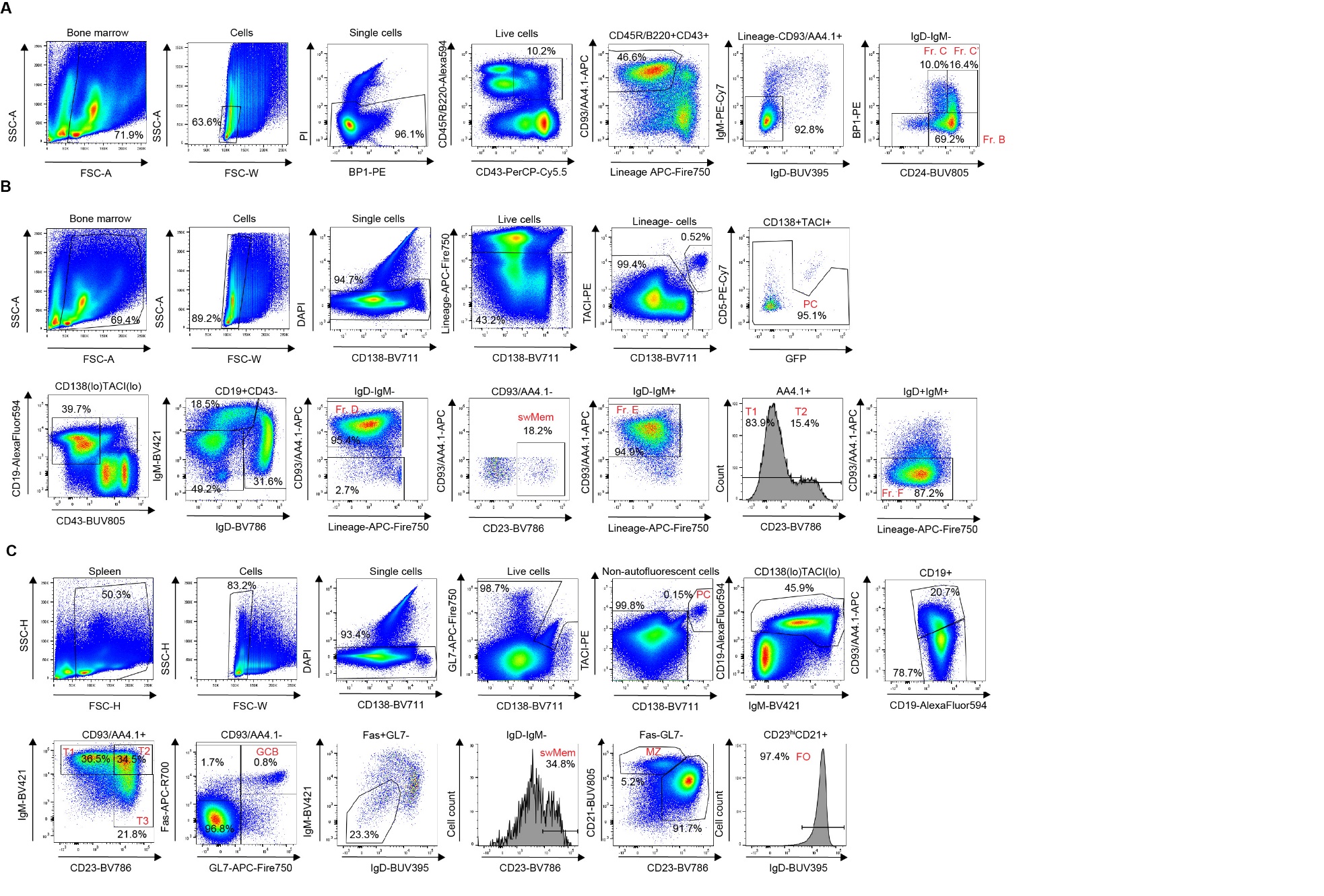

**Supplementary Fig. 1: Gating strategy for B cell subset identification.**

**(A–C)** Mice aged 9–13 weeks were analyzed by flow cytometry. Representative pseudocolor plots and histograms from a wild-type control mouse illustrate the gating strategy used to identify B cell subsets in **(A, B)** bone marrow and **(C)** spleen.

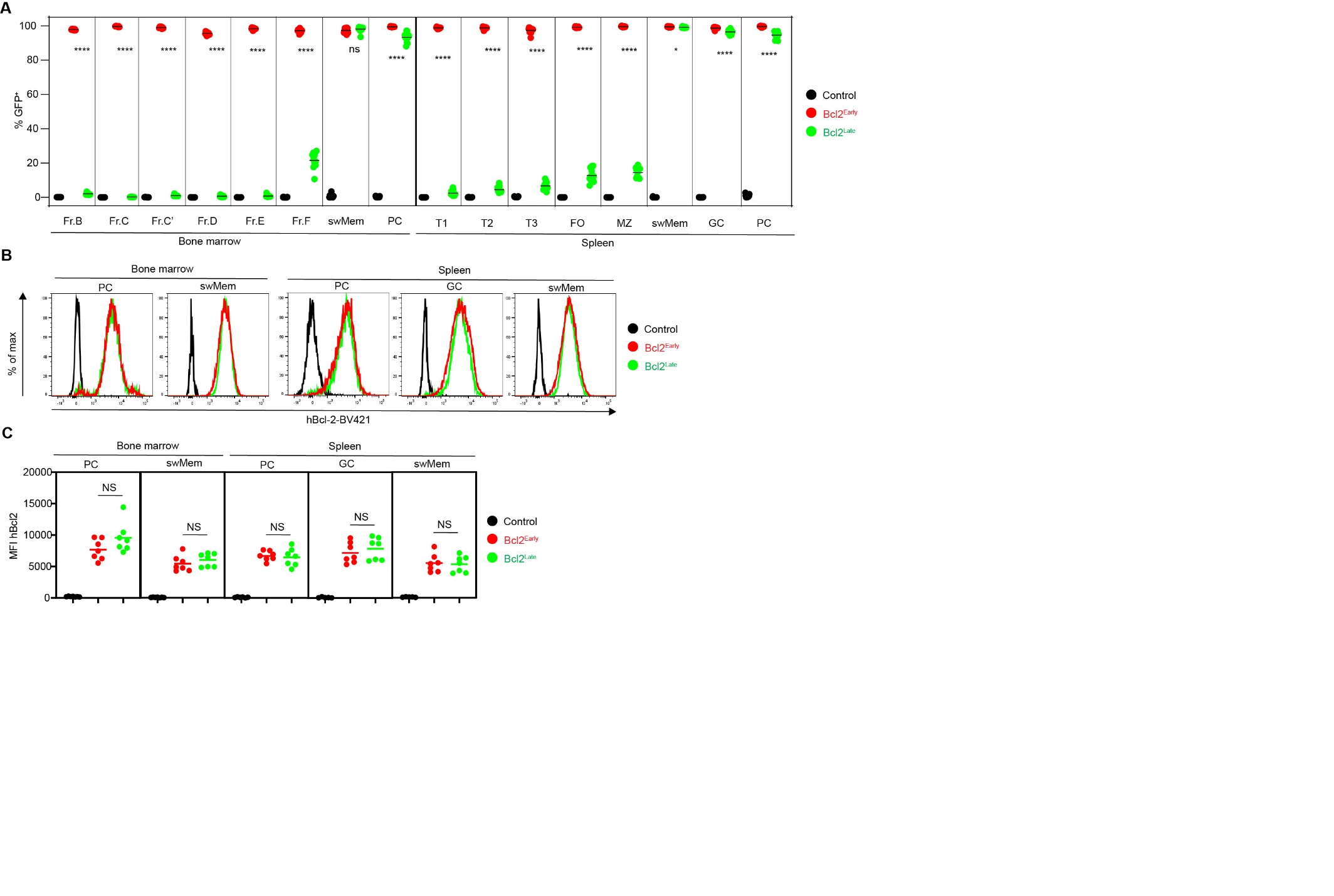

**Supplementary Fig. 2: Validation of GFP and hBcl-2 expression in Bcl2^Early^ and Bcl2^Late^ mice.** (**A-C**) Mice aged 9-13 weeks were analyzed by flow cytometry. (**A**) Quantification of GFP expression among indicated B cell subsets in Control (black), Bcl2^Early^ (red) and Bcl2^Late^ mice (green). GFP is co-expressed with hBcl-2 following Cre-mediated recombination. Data are combined from two to six independent experiments (Controls, n=16; Bcl2^Early^, n=11; Bcl2^Late^, n=12). (**B, C**) Mice aged 12-20 weeks were analyzed for intracellular human Bcl-2 (hBcl-2) expression by flow cytometry. (**B**) Representative histograms show hBcl-2 expression across indicated B cell subsets and genotypes. (**C**) Quantification of hBcl-2 mean fluorescence intensity (MFI). Data are combined from two independent experiments (n=7 per group). **** p<0.0001, *** p<0.001, ** p<0.01, * p <0.05, NS=not statistically significant (two-tailed Mann-Whitney test for A; two-tailed unpaired Student’s t-test for C). Horizontal bars indicate mean values. Abbreviations: swMem, class-switched memory B cells; PC, plasma cells; T1, transitional 1 B cells; T2, transitional 2 B cells; Anergic/T3, anergic B cells; FO, mature follicular B cells; MZ, marginal zone B cells; GC, germinal center B cells.

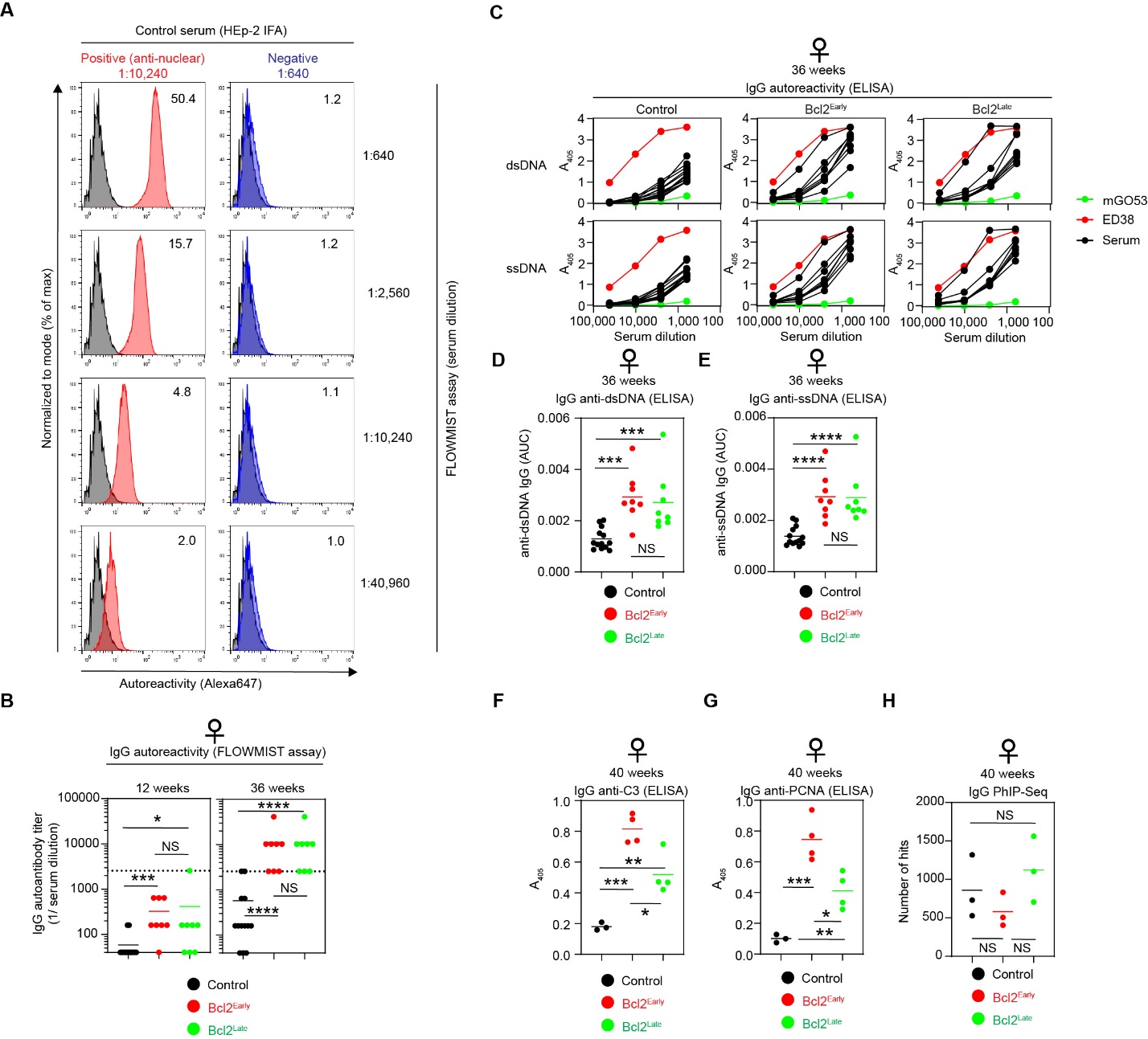

**Supplementary Fig. 3. Serum IgG autoantibody reactivity**.

**(A**) Validation of the FLOWMIST assay used to quantify autoreactive IgG in mouse serum. Representative histograms show intracellular staining of splenocytes with indicated dilutions of control sera previously characterized by HEp-2 cell immunofluorescence (HEp-2 titer shown above), followed by Alexa647-conjugated secondary antibody (red, positive control with anti-nuclear antibody titer 1:10,240; blue, negative control with no HEp-2 reactivity at 1:640 dilution; grey, unstained control). Numbers indicate mean fluorescence intensity (MFI) ratios of stained versus unstained samples; ratios above 3 were considered positive. **(B)** Serum IgG autoantibody titers measured by FLOWMIST in 12- or 36-week-old female mice of the indicated genotypes. Values at or above the dotted line indicate high-titer autoreactive IgG (≥ 1:2560). **(C-E)** Serum IgG binding to double-stranded DNA (dsDNA) or single-stranded DNA (ssDNA) measured by ELISA. (**C**) PBS-corrected absorbance (A_405_) values plotted as a function of serum dilution. ED38 is included as a highly polyreactive control antibody and mGO53 as a non-reactive control antibody. (**D, E**) Quantification of binding to (**D**) dsDNA or (**E**) ssDNA expressed as area under the curve (AUC). (**F, G**) Quantification of serum IgG binding to (**F**) complement C3 or (**G**) PCNA measured by ELISA. (**H**) Serum IgG autoreactivity assessed by phage immunoprecipitation sequencing (PhIP-seq). Symbols indicate the total number of significant peptide hits detected per mouse, grouped by genotype. (**B, D, E**) **** p<0.0001, *** p<0.001, * p <0.05, NS=not statistically significant (two-tailed Mann-Whitney test). (**F-H**) *** p<0.001, ** p<0.01, * p<0.05, NS, not statistically significant (two-tailed unpaired Student’s t-test).

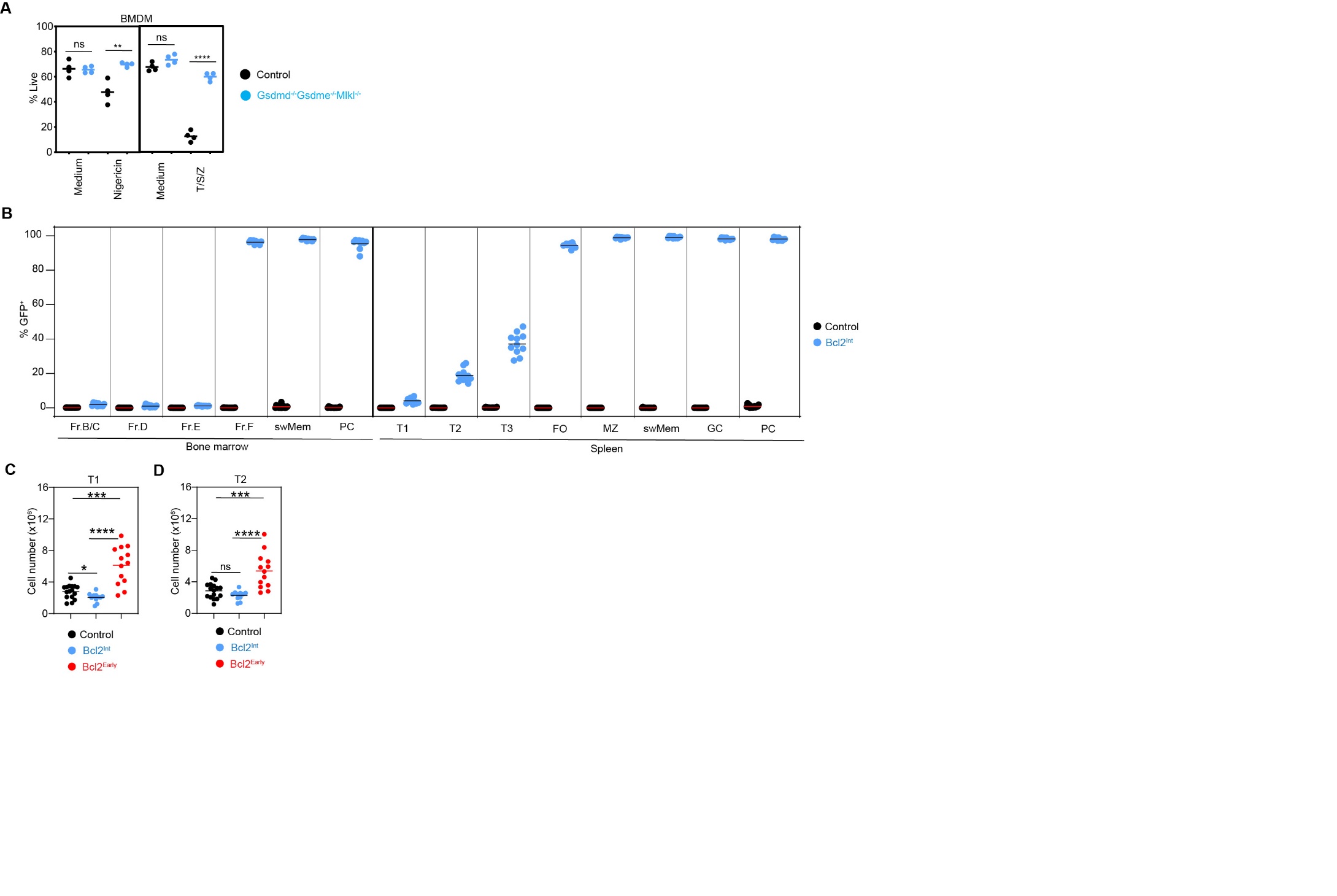

**Supplementary Fig. 4. Control experiments and validation of Bcl2^Int^ mice**. (**A**) Pam3CSK4-primed bone marrow derived macrophages (BMDM) were stimulated with nigericin for 1h to trigger pyroptosis or with TNFα (T), birinapant (SMAC mimetic; S) and Z-VAD-FMK (Z) (T/S/Z) for 16h to induce necroptosis. Following propidium iodide staining, viability was assessed by flow cytometry. One representative experiment out of three independent experiments is shown (n=4 per group). (**B-D**) Mice aged 9-13 weeks were analyzed by flow cytometry to assess CD21^Cre^Rosa26^LSL-Bcl2^ (Bcl2^Int^) recombination and developmental effects. Bcl2^Int^ mice were analyzed in parallel with Control and Bcl2^Early^ cohorts shown in (fig. S2A-C). (**B**) Quantification of GFP expression among indicated B cell subsets. (**C, D**) Quantification of total numbers of (**C**) T1 and (**D**) T2 cells in the spleen (Controls, n=16; Bcl2^Early^, n=13; Bcl2^Int^, n=11). Data are combined from two to six independent experiments. **** p<0.0001, *** p<0.001, ** p<0.01, * p <0.05, NS=not statistically significant (two-tailed unpaired Student’s t-test for A; two-tailed Mann-Whitney test for C and D). Horizontal bars indicate mean values. Abbreviations: swMem, class-switched memory B cells; PC, plasma cells; T1, transitional 1 B cells; T2, transitional 2 B cells; Anergic/T3, anergic B cells; FO, mature follicular B cells; MZ, marginal zone B cells; GC, germinal center B cells.
